# Deep Learning Features Encode Interpretable Morphologies within Histological Images

**DOI:** 10.1101/2021.08.16.456518

**Authors:** Ali Foroughi Pour, Brian White, Jonghanne Park, Todd B. Sheridan, Jeffrey H. Chuang

## Abstract

Convolutional neural networks (CNNs) are revolutionizing digital pathology by enabling machine learning-based classification of a variety of phenotypes from hematoxylin and eosin (H&E) whole slide images (WSIs), but the interpretation of CNNs remains difficult. Most studies have considered interpretability in a post hoc fashion, e.g. by presenting example regions with strongly predicted class labels. However, such an approach does not explain the biological features that contribute to correct predictions. To address this problem, here we investigate the interpretability of H&E-derived CNN features (the feature weights in the final layer of a transfer-learning-based architecture), which we show can be construed as abstract morphological genes (“mones”) with strong independent associations to biological phenotypes. We observe that many mones are specific to individual cancer types, while others are found in multiple cancers especially from related tissue types. We also observe that mone-mone correlations are strong and robustly preserved across related cancers. Importantly, linear mone-based classifiers can very accurately separate 38 distinct classes (19 tumor types and their adjacent normals, AUC=97.1% ± 2.8% for each class prediction), and linear classifiers are also highly effective for universal tumor detection (AUC=99.2% ± 0.12%). This linearity provides evidence that individual mones or correlated mone clusters may be associated with interpretable histopathological features or other patient characteristics. In particular, the statistical similarity of mones to gene expression values allows integrative mone analysis via expression-based bioinformatics approaches. We observe strong correlations between individual mones and individual gene expression values, notably mones associated with collagen gene expression in ovarian cancer. Mone-expression comparisons also indicate that immunoglobulin expression can be identified using mones in colon adenocarcinoma and that immune activity can be identified across multiple cancer types, and we verify these findings by expert histopathological review. Our work demonstrates that mones provide a morphological H&E decomposition that can be effectively associated with diverse phenotypes, analogous to the interpretability of transcription via gene expression values.

## 1. Introduction

Deep learning has become an important methodology for analyzing biomedical images, and in particular for analyzing hematoxylin and eosin (H&E) stained whole slide images (WSIs). Deep neural networks have achieved classification accuracies higher than classical machine learning models [1]. However, they are black-boxes that do not directly reveal the morphological features they associate with labels, a significant concern for mechanistic analysis and clinical decision making [2]. Identification of biologically meaningful morphological features may be confounded by image artifacts [3], such as blurring, noise, and lossy image compression [4]. Tissue damage, image quality, and dataset-specific artifacts have also been suggested to affect feature representation and prediction accuracy of neural networks (see [1], [5], and [6]). Given the impact of such artifacts on deep learning-based predictors, it is of critical importance to be able to decompose CNNs into features that can be biologically interpreted.

The majority of models for visualizing, analyzing, and interpreting CNNs reveal “where” a network is “looking” to make its prediction, rather than revealing “what” information in the region of interest is important. Some methods output pixel patterns that affect the value of a neuron in a deep network (see [7]). However, such techniques tend to output different predictive regions, can be difficult to validate, or have been suggested to be “fragile”, i.e. extremely sensitive to small perturbations of the image [8]. Optimizing conventional deep learning techniques to identify regions informative of class labels is a current theme in digital pathology (see [9, 10] for examples). Other recent works have focused on visualizing individual deep learning features as heatmaps [11].

Unlike natural image analysis [12], biomedical image analysis is complemented by additional data modalities, such as multiplexed imaging, single cell and bulk sequencing, and clinical information citep ray2014information, kong2011integrative. These data may aid in interpreting the deep feature representations of the H&E slide. However, models integrating these diverse modalities are needed. The feasibility of doing so is supported by work establishing the connection between modalities, for example by using CNNs to predict expression values of specific genes from H&E images (see [13, 14, 15] for examples). Because of the architectural complexity of CNNs, it has often been assumed that CNN-based decompositions of images into features are not interpretable. However, there has been little empirical study of this question, e.g. by testing whether CNN-derived features are correlated with simple biological features such as gene expression values.

In this work, we investigate the interpretability of CNN-derived image features, which we conceptualize as morphological genes (“mones”). We find that mones have strong linear associations with phenotypic features, making them directly interpretable, which we demonstrate in several analyses. We demonstrate that many mones can distinguish cancer tumors from adjacent normal slides. These mones can be linearly combined for reliable prediction of both pan-cancer and tissue-specific cancer phenotypes. Mone-mone correlation analysis identifies clusters of highly correlated mones within cancer types, and these correlations are strongly preserved among cancers from related tissues. Mone values resemble gene expression data, allowing immediate use of many interpretable bioinformatics tools and machine learning models to identify the underlying biology of morphologies encoded by CNNs. For example, integrative mone-gene expression correlation analysis reveals that collagen content and immune infiltration are linearly associated with morphologies encoded by mones in several cancer types, and we confirm these relationships by expert histopathological review. Our studies demonstrate the power of mones for computationally deconstructing cancer images into interpretable biological features.

## 2. Results

We analyzed the InceptionV3 [16] features of tiles from whole slide images of The Cancer Genome Atlas (TCGA) [1] for 19 cancers (see Supplementary File 1 for the full list). We used features derived from this architecture because predictive models based on Inception have shown high accuracy for identifying phenotypes in prior studies [1]. We hereafter denote each of the 2048 outputs of the global average pooling layer of the Inception V3 network as mones (morphological genes). We use this terminology because mones have analogies to genes with individual expression values. Tile level mones can be combined to construct slide level mones (see methods). Unless otherwise stated, in the studies below “mone” refers to a slide level characterization.

### 2.1. Individual mones differentiate phenotypes

We first investigated to what extent individual mones can differentiate phenotypes, focusing on TCGA tumor/normal slide comparisons. We initially identified individual mones with significant differences in distribution between breast cancer (BRCA) tumor and adjacent normal slides (>1800 mones were statistically significant with FDR< 5%). A clustermap of the top 100 such mones was able to clearly separate these classes (see methods, Figure 1a, and Supplementary Figure 1, clustermap AUC=89%, rand score =96%, and adjusted rand score=85%). These results were typical of mone behaviors in many tumor types – in any given tumor type, many mones were able to separate frozen tumor from normal slides (see methods and Supplementary File 1). We applied several statistical methods to test the robustness of such mones. In each cancer, at least 40% of mones were statistically significant irrespective of the statistical test used (FDR= 5%, see Supplementary File 1), and 75.7% of mones significant by at least one method were significant by all tests (see methods, see Figure 1b). A smaller subset of these mones showed strong effect sizes, as identified by optimal Bayesian filter (OBF) [17] statistics (see Methods). 22% of all mone-cancer pairs met this criterion based on distributional differences between tumor and normal slides (see Supplementary Figure 2).

**Figure 1:**
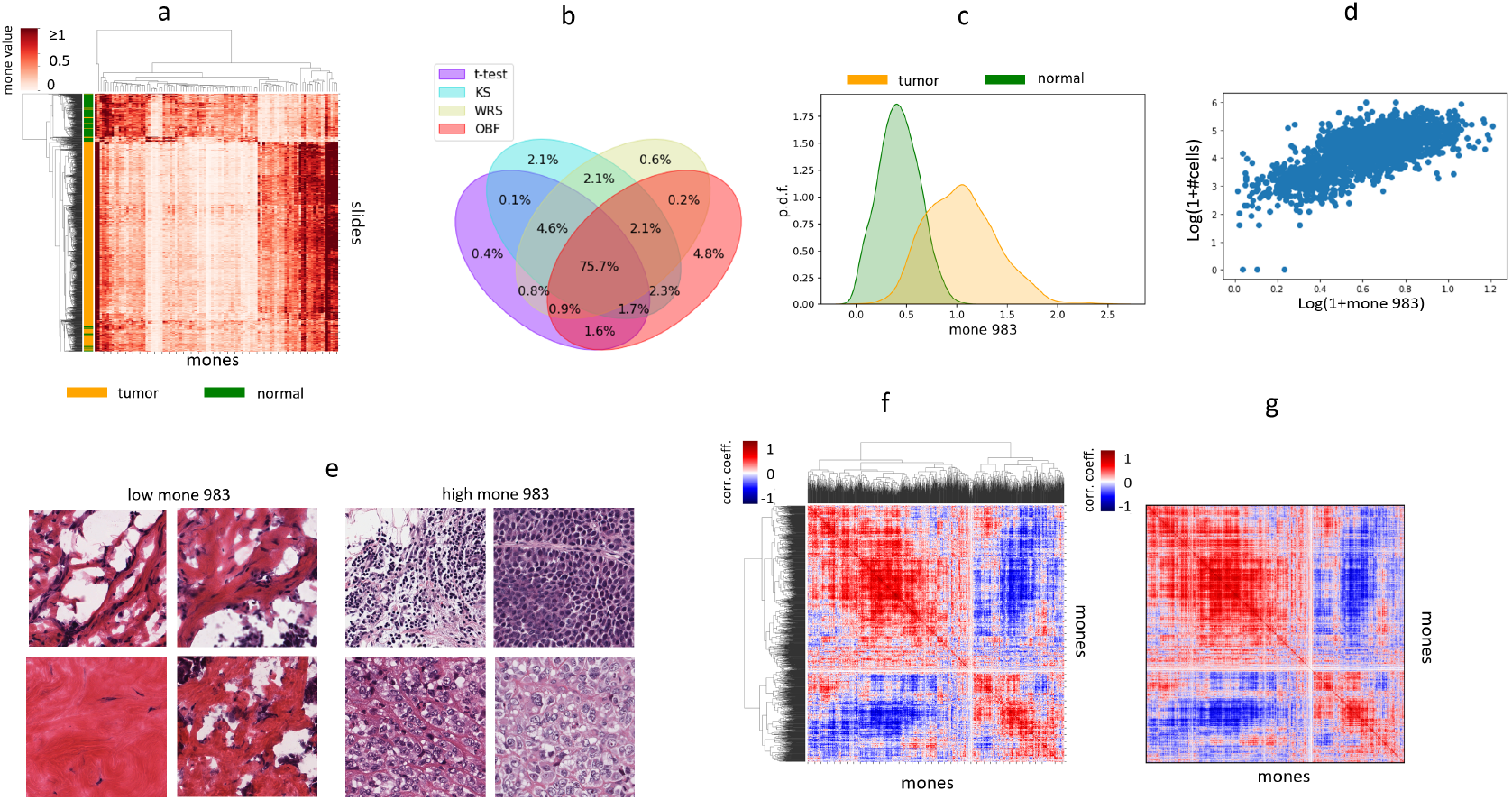
Individual mones and mone pairs encode and distinguish phenotypes. (a) Clustermap of BRCA slides using the top100 mones differentiating the slides. 100 mones are sufficient to separate frozen normal (green) from frozen tumor (orange) slides. (b) Venn diagram of statistically significant mones differentiating tumor from adjacent normal frozen slides, comparing different statistical tests. Venn diagrams were calculated for each cancer type, and the observed plot shows the average across all cancer types. On average the statistical tests agree on 75% of mones differentiating between tumor and normal slides. (c) Probability density function of mone 983 among frozen tumor (orange) and adjacent normal (green) BRCA slides. (d) Log-normalized scatter plot of slide level mone 983 and cellpose (see [48]) estimates of cellularity across BRCA frozen slides. (e) Example tiles from slides with extreme mone 983 values (high and low). (f) Cluster map of the mone-mone correlation matrix of LUAD tumor slides, demonstrating that many mones pairs are highly correlated. (g) Mone-mone correlation matrix of LUSC slides, with mones ordered identically to the Figure 1f cluster map.

As an example, mone 983 is a tumor marker with a distributional difference between frozen tumor and adjacent normal breast cancer (BRCA) slides (Figure 1c). It also behaves similarly in several other tumor types (Supplementary Figure 3). It strongly correlates with cell density in frozen BRCA slides (see methods, see Figures 1d-e, Pearson r=0.69, p-value<1e-200, see Supplementary File 2) and significantly though with moderate magnitude in FFPE BRCA tumor slides (Pearson r=0.24, p-value=1.9e-13). Mone 983 has higher correlation with cell density in FFPE LUAD slides (Pearson r=0.61, p-value=2.2e-34) than frozen LUAD slides (Pearson r=0.28, p-value=4.1e-18, see Supplementary File 2).

To further test whether mones exhibit consistent behavior in different cancer types, we analyzed the four cancer families of [1]: pan-GYN, pan-KIDNEY, pan-LUNG, and pan-GI (see Supplementary File 3 for cancers in each family). More than half of the mones with distributional differences in each cancer type also have distributional differences in all cancers of the family (see Supplementary File 3). Although cancer-specific mones are uncommon, such mones still show clear distributional differences between tumor and normal (see Supplementary Figure 4). Interestingly, most mones distinguishing tumor from normal in both LUAD and LUSC also distinguish between the LUAD and LUSC cancers (see Supplementary File 3), suggesting quantitative values are important. Individual mones can also distinguish frozen from FFPE slides (see Supplementary File 4). We observed the ability of some mones to distinguish between tumor and normal is impacted by differences between frozen and FFPE modalities (421 ± 112 across all cancers, see methods, see Supplementary File 4), though the majority of mones behave similarly in frozen and FFPE.

### 2.2. Mone clusters provide robust encodings of cancer phenotypes

We next investigated to what extent mones are independent or encode behaviors together. We did this by calculating pairs of mones that were significantly correlated, for each cancer type. We analyzed this first by restricting to frozen slides (to avoid Simpson’s paradox), and second by combining frozen and FFPE slides (to avoid false correlations due to frozen-specific artifacts). The results were generally robust between the two methods—a few cancers had a non-trivial difference in the ratio of correlated mone pairs ascertained by the two methods (ESCA, KICH, SARC, and PRAD 22.3% ± 2.22%), while the remaining cancers had small differences (7.8% ± 3.9%, see Supplementary File 6). Therefore we subsequently analyzed frozen samples only unless otherwise specified. Overall, we found that correlated mones are prevalent among tumor slides (see Figures 1f and 1g, and Supplementary File 6). For example, 83.9% and 88.7% of mone-mone pairs are correlated within LUAD and within LUSC, respectively (see methods and Supplementary File 6). We observed similar results for other cancers: 68.8% ± 13% of all mone-pairs were statistically significant across cancers (see Supplementary File 6). Remarkably, we observed that pairwise correlations are preserved within cancer families – more than 45% of mone-pairs statistically significant in one cancer are significant in all cancers of the family (pan-GYN, pan-KIDNEY, pan-LUNG, and pan-GI, see Supplementary Figure 5). For example, the mone-mone correlations in lung adenocarcinoma and lung squamous cell carcinoma are nearly identical (Figures 1f and 1g). We also calculated mone-mone correlations in the normal slides associated with each cancer type, hypothesizing that the difference in mone-mone correlations between tumor and normal might be important to distinguishing tumor from normal images. However, these differential mone correlations are weaker and less preserved across cancers (see Supplementary File 6 and Supplementary Figure 5).

Different cancers can be distinguished by different sets of mones. For example, while mone 983 separates tumor and normal slides of BRCA, it does not differentiate COAD tumor and normal slides (see Supplementary Figure 3). We identified 105 mones that are highly correlated with mone 983 in BRCA (Null: |*r*| *≤* 0.5, FDR= 0.1%, see methods), among which 52, 14, and 29 also differentiate tumor from normal slides in COAD, READ, and STAD, respectively (t-test FDR<0.1%). The first principal component of these COAD-overlapping mones (explaining 41% of the variance across COAD samples) strongly correlates with cell density in frozen slides (Cellpose cellularity estimates: Pearson r=0.22, p-value=2.2e-11, HoverNet cellularity estimates: Pearson r=0.43, P-value=1.9e-47, see Methods and Supplementary File 2). Thus the high cellularity in BRCA [18, 19] and COAD [20, 21] involve incompletely overlapping mone sets.

### 2.3. Linear models of mones can detect and distinguish tumors

We investigated linear models of mones for predicting phenotypes, as they allow direct interpretation of mone values. We observed that linear models of mones can efficiently distinguish tumor from adjacent normal slides, as well as the cancer type from which they are derived (19 cancers, 38 classes, see methods, see Figure 2a-c). We tested two linear models, multiclass linear discriminant analysis [MLDA, One versus Rest (OVR)-AUC=97.1% ± 4.6%, see Supplementary File 7] and multinomial logistic regression with LASSO penalty (LR-LASSO, OVR-AUC=97.1% ± 4.2%, see Supplementary File 7). MLDA encodes mone patterns indicative of class labels into a low dimensional space (i.e. the number of classes - 1), yielding t-SNE visualizations with improved interpretability over naïve t-SNE (compare Figures 2a,b and Supplementary Figures 6a,b). LR-LASSO, on the other hand, is a linear model based on small mone sets, so its regression coefficients can be interpreted directly with risks incurred by each mone. Although CNN methods typically use difficult-to-interpret fully connected layers at the classification step, we found that efficiently designed linear models can replace fully connected layers while still achieving high prediction AUCs. Combining tumor probabilities of the LR-LASSO classifier we obtained a universal tumor detector with extremely high AUC (99.2% ± 0.12%, see methods), out-performing the fully deep learning model of [22] (Reported AUC=0.95 ± 0.02). Furthermore, LR-LASSO is effective at cross-classification similar to CNNs with fully connected classification layers [1], i.e., LR-LASSO trained to distinguish tumor/normal for one cancer type can distinguish tumor/normal for other cancer types as well (Figure 2d and Supplementary Figure 6c). While the LR-LASSO model has smaller average AUC (0.84) compared to the fully deep learning model of [1] (0.88), logistic regression is more interpretable than a multi-layer perceptron. Our slide level tumor detectors also produce meaningful tile level predictions. Independent review by our pathology team supports most tumor regions having high tumor probability, and most non-tumor regions have low tumor probability in these images, with the cases of misclassification tending to be prediction of non-tumor regions to be tumor (see Supplementary Figure 7 for examples). These results indicate that the LR-LASSO slide level tumor markers are effective at the tile level.

**Figure 2:**
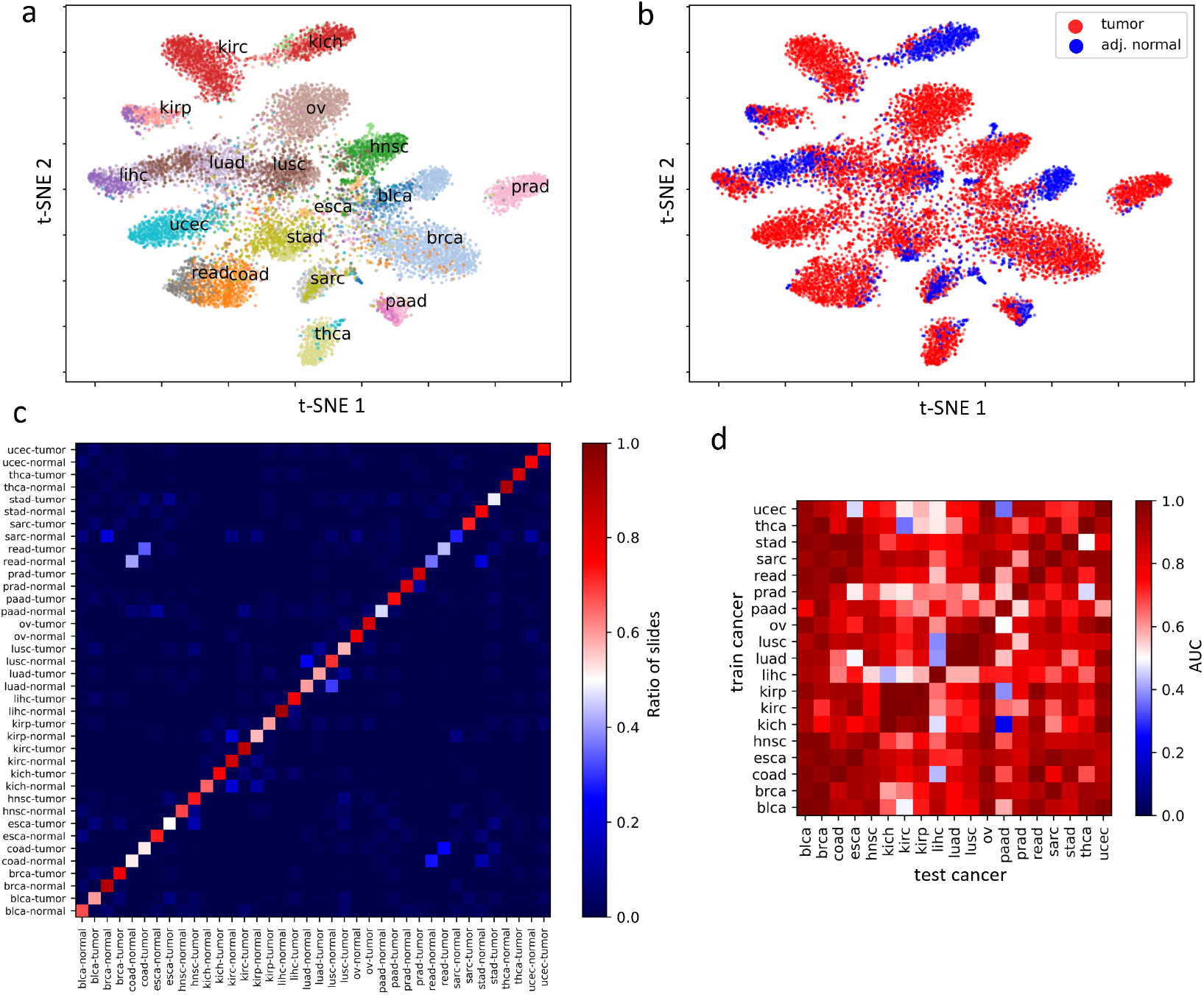
The joint distribution of mones reliably separates tumor and normal slides and the underlying cancer. 2D t-SNE plots of the mone-based MLDA feature space distinguishing 38 classes (19 cancers, tumor/normal status) based on (a) cancer type and (b) tumor/normal status. (c) Normalized confusion matrix of the 38-class mone-based logistic regression classifier. The color depicts the ratio of slides with a given true class predicted as any of the possible classes. The large diagonal values suggest the classifier has high accuracy. (d) The cross-classification AUCs of mone-based logistic regression tumor/normal classifier trained on each cancer and applied to all cancers.

### 2.4. Mones have interpretable correlations with gene expression

We next investigated whether mones are linearly associated with gene expression values, as this could provide transcriptional interpretability for mones (see Methods for data stratification and pre-processing). We observed many mones significantly correlated with individual genes. Across five cancers analyzed (OV, COAD, KIRC, LUAD, and LUSC) between 83 (LUSC) and 1797 (KIRC) mones were associated with at least one gene, whereas between 332 (LUSC) and 16474 (KIRC) genes were associated with at least one mone (see methods, see Supplementary File 8). We then analyzed several cases of particular interest.

#### 2.4.1. Mones encode collagen content

We first used unsupervised analysis to study clusters of correlated mone-gene pairs. We identified a cluster of highly correlated mones and collagen genes in OV (Figure 3a). These mone values can be efficiently combined for association with phenotype using PCA (PC-1 explains 63% of the variance). Histopathological review by our pathology team confirmed tiles with high PC-1 values as typically rich in collagen (see Figure 3d and Supplementary Figure 8), and tiles with low PC-1 values as having low collagen but increased cellularity. These mone-gene associations may be clinically relevant, as high expression of collagen genes correlates with multi-drug resistance [23] and poor prognosis in ovarian cancers [24]. Mone 1062–one of the mones in the identified cluster– was additionally highly correlated with ECM2, THBS1 and THBS2, which have been suggested to play a significant role in ovarian cancer drug resistance and metastasis [24, 25, 26].

**Figure 3:**
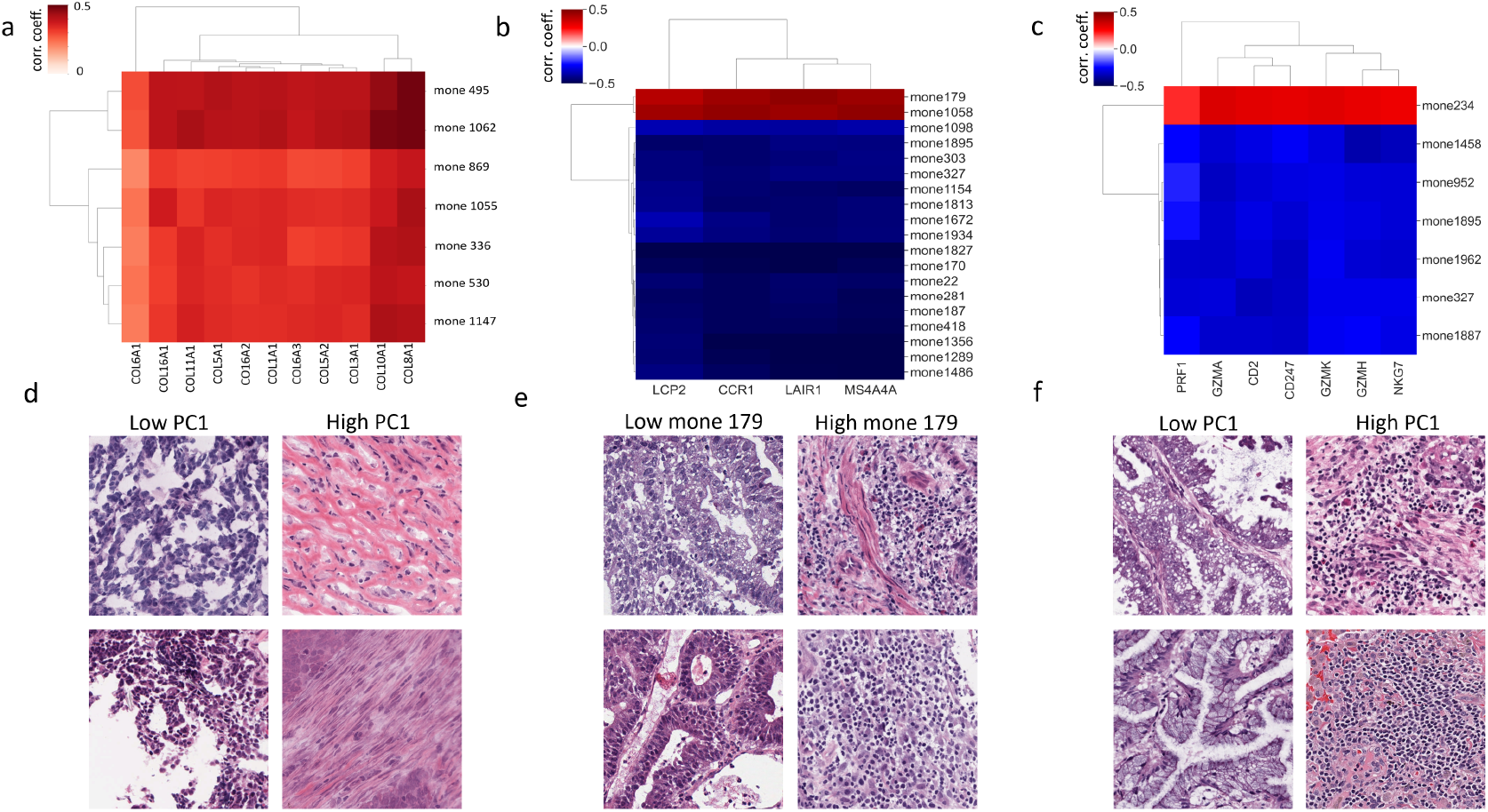
Mone-gene correlation analysis identifies highly correlated mone-gene clusters. Correlation matrix of (a) a cluster of highly correlated mones and collagen genes in OV, and a cluster of highly correlated mones and immune-related genes in (b) COAD and (c) LUAD. See Supplementary Figure 8 for adjusted p-values. Example tiles from slides with (d) high and low PC-1 in OV, (e) high and low mone 179 in COAD, and (f) high and low PC-1 in LUAD. Histopathology review identifies that mone-predicted (d) OV tiles with high PC-1 are rich in collagen, and (e) COAD with high mone 179 and (f) luad tiles with high PC-1 have a strong lymphocyte presence. See Supplementary Figures 9-12 for additional examples at both high and low mone values.

#### 2.4.2. Mones encode immune infiltration

Supervised correlation analysis using fixed gene sets can also be used to test if mones encode morphological features associated with a biological phenotype. We tested whether mones encode immune infiltration in pan-GI (COAD, READ, and STAD) and pan-LUNG (LUAD and LUSC) cancers, with immune related gene sets taken from [27] and [28] (see Methods and Supplementary File 9).

We identified 19 mones significantly correlating (some positive, some negative) with immune related genes in pan-GI cancers (Fig 3b and Supplementary File 10). These mones have significant correlations with each other (|*r*|= 0.56 ± 0.13), and 14 mones significantly correlate with more than one immune gene. LAIR1, LCP2, MS4A4A, and CCR1 correlate with 15, 9, 12, and 15 mones, respectively. All 19 mones are significantly correlated with prior TCGA estimates of leukocyte fraction [29] (FDR<0.05,|r|=0.29 ± 0.07) and HoverNet estimates of immune cell quantity from the H&E images (see methods, FDR<0.05, |*r*|= 0.45 ± 0.12) in COAD. Mone 179 had strong positive correlations with both leukocyte fraction (*r* = 0.2) and HoverNet estimates (*r* = 0.65). Histopathological review confirmed that mone 179 differentiates amongst COAD tiles according to their level of immune infiltration (see Figure 3e and Supplementary Figure 9). PC-1 of the 19 mones has significant correlation with leukocyte fraction (*r* = 0.4, *p — value* = 2.4*e -* 35) and HoverNet estimates (r=0.17, p-value=2.5e-17). Histopathological review validated that PC-1 also differentiates COAD tiles according to differing levels of immune infiltration (see Supplementary Figure 10). Thus PC-1 efficiently combines mones and provides a stronger separation than individual mones.

We observed similar correlations, but smaller in magnitude, between mones and local immune cytolytic activity genes in lung cancers (Fig 3c and Supplementary File 10). We observed stronger correlations in LUAD than LUSC (LUAD |r|=0.20 ± 0.05, LUSC |r|=0.12 ± 0.05). We identified 31 mones significantly correlating (some positive, some negative) with immune related genes in LUAD. 7 mones correlated with at least 3 genes (see Figure 3c). Only one LUAD immune mone did not correlate with lymphocyte fraction (FDR=0.05). PC-1 of the 31 mones correlates with lymphocyte fraction (r=0.28, p-value= 2.2e-4). Histopathological review of LUAD slides based on PC-1 suggests LUAD tiles with high PC-1 show a strong tumor infiltrating lymphocyte presence and have inflammation, while tiles with low PC-1 are typically not inflamed and show weaker immune infiltration (Fig 3f, Supplementary Figure 11).

#### 2.4.3. Mones identify immunoglobulin gene expression in highly cellular colon adenocarcinoma tumors

Supervised correlation analysis can also clarify finer behaviors within WSIs. For example, highly cellular COAD tumors typically show high expression of immunoglobulin (IG) genes. We considered the 52 mones correlated with mone 983 and which differentiate COAD tumor from normal (see section 2.2). The PC-1 dimension of these mones significantly correlates with 87 genes in COAD (FDR<0.05, see Supplementary File 11 for the full gene list), as well with the average expression of these 87 genes (r=0.33, p-value=3.8e-12). Immunoglobulin (IG) genes dominate this gene set, and we define their average expression as a sample’s IG score. B-cells express immunoglobulin, and as expected we observed a statistically significant correlation between the log normalized B-cell estimates of [29] and IG score (p-value=2.4e-12). Interestingly, however, B cells comprise only a small fraction of cells in each sample (0.86% ± 1.5%), and we did not observe a significant correlation between the B-cell estimates and mone PC-1 (correlation coefficient=-0.009, P-value=0.85). Thus mone analysis suggests that IG expression is not due solely to B-cells. This supports recent studies suggesting colon cancer cells themselves express IG genes (see [30]).

## 3. Discussion

Mones enable interpretation of deep neural networks because they linearly correlate with phenotypes and gene expression profiles. We have empirically shown that mones efficiently encode strong morphological features that can often be used to replace multi-layer perceptrons with robust and interpretable linear classification models. Moreover, we have demonstrated that integrative mone-gene correlation analysis can identify specific transcriptional processes from images, and verified these through expert pathological review.

### 3.1. Mones provide image-based interpretability

Linear models based on mones have several empirical and theoretical strengths for image analysis. Individual mones and small mone clusters directly correlate with phenotypes (see section 2.1 and 2.3), enabling a simpler interpretation of CNNs compared with methods that integrate all CNN features together. Mones decompose arbitrary images into interpretable features, in contrast to approaches that assume non-linearity and which therefore provide interpretability primarily through example regions identified by problem-specific classifiers. A notable advantage of the mone approach is that it does not require a trained classifier.

Some prior linear analyses of deep learning features have enabled partial interpretation of CNNs trained on one cancer type (see [11] and [31]), but mones facilitate pan-cancer interpretations. For example, our results demonstrate a pan-cancer mone can encode conserved morphological features across multiple cancer types (see section 2.1), and a conserved morphological feature can be encoded by distinct mones in different cancer types (see section 2.2). Moreover, mones within strongly correlated clusters can be combined to better identify shared encoded morphology (see section 2.2 and [32]).

Mone analysis improves low dimensional visualization of large image sets, such as via t-SNE plots. Pre-trained CNNs are universal feature extractors that encode morphological features predictive of a multitude of labels (see [1], [33], and [34]). Because not all high-dimensional complex relations can be easily embedded in 2D, mone analysis can be used as a first phase of targeted search to identify the mones relevant to classes of interest (see section 2.3), allowing more accurate visualization of class separations in mone space.

Correlation analysis of mones with gene expression values is a powerful approach for interpreting mones. We identified clusters of highly correlated mone-gene sets, demonstrating clear connection of mones to the underlying genetics. Some recent studies have used exhaustive sets of deep learning features to predict expression profiles (see [13, 14, 15]), but our work shows that small mone-gene clusters can be sufficient and provide simpler interpretability. Both supervised and unsupervised analyses identify meaningful clusters (see section 2.4). Supervised analysis using fixed gene sets is particularly interesting, as it enables direct assessment of genes of interest via mones (see section 2.4.2).

### 3.2. Mone analysis across architectures and data modalities

Our analyses demonstrate that mones provide an efficient and interpretable CNN embedding of image data, but a caveat is that they have been restricted to Inception v3 mones. Architectures with fully connected layers tend to increase non-linearity in feature representations. Therefore, models that do not utilize multiple sequences of fully connected layers, such as Inception, are more appropriate for linear mone analysis. For example, recent work suggests a small subset of VGG19 features may also be interpreted via their direct association with phenotypes [11]. However, we believe Inception V3 mones are more appropriate for linear association studies because they are the direct inputs to the classification layer. A few other studies have explored correlations between deep autoencoder features and gene (see [31]) or protein expression [35] for other architectures, but the relations between mones across architectures remains a broad and open research topic. We have found that Inception V3 mone 983 in BRCA can be reliably estimated via linear models using ResNet152V3 and DenseNet201 mones (*R*^2^ > 0.95). Furthermore, we have been able to convert Xception mones to Inception V3 mones using autoencoders with reasonable accuracy (*R*^2^ ≈ 0.5). These studies suggest that integrative linear mone-gene correlation analysis can be made effective across a range of deep learning architectures.

While this work focuses solely on H&E WSIs, we believe mone-based interpretation will be valuable for extension to other spatial data types. Immunohistochemistry (IHC) images are primed for rapid progress, as recent work has shown that several IHC markers can be virtualized from H&E (see [36] and [37]). Generalizing mone analysis for other data types such as spatial transcriptomics and multi-channel protein data is also an exciting and open area, though new architectures will need to be explored to handle the high dimensionality of such images. The interpretation of CNNs for these image types is a challenging but important task, and we expect that integrative multi-modal multi-architecture mone analysis will be a potent and informative approach.

## 4. Methods

### 4.1. Data acquisition and pre-processing

The 20X H&E WSIs of TCGA are pre-processed, tiled, and passed through an InceptionV3 model pre-trained on image-net data in [1]. The cached 2048 global average pooling layer features of InceptionV3 (called mones in this manuscript) were written to disk and analyzed. For each of these 2048 mones, the median value across all tiles within a slide was computed to yield the slide-level mone value.

### 4.2. Differential mone analysis

Differential mone analysis identifies mones with statistically significant distributional differences across classes. Welch’s t-test, Kolmogorov Smirnov (KS) test, Wilcoxon Rank Sum (WRS) test, and optimal Bayesian Filter (OBF) (see [17] for details) were used for statistical analysis. t-test, WRS test, and KS test use the Benjamini-Hochberg procedure [38] for FDR correction. The scipy python package [39] was used to implement t-test, KS test, and WRS test. The statsmodels [40] implementation of the Benjamini-Hochberg procedure was used.

Minimal risk OBF (see [17] for details) identifies mones with posterior probabilities larger than 1 − *α*, where *α* is the FDR rate. FDR-OBF (see [41] for details) outputs the feature set that bounds the sample conditioned FDR by *α*. OBF can report the FDR of any arbitrary feature set. Unless otherwise stated FDR-OBF is used. OBF uses Jeffrey’s prior, assumes the prior probability of a mone having distributional differences is 50% (to model no preference on the identity of a mone, i.e., with or without distributional differences across classes), and sets the normalization constant of the prior to 0.1. Mones with distributional differences across classes are hereafter called markers, and mones without distributional differences are called non-markers. The posterior probabilities of OBF can be used to estimate the first two moments (mean and standard deviation) of the number of markers (see [41] for details). Assuming mone identities (marker or non-marker) are independent across cancers, the posterior probabilities can be multiplied to calculate the probability of a joint event.

OBF intrinsically computes the ratio between sample variance and weighted geometric mean of class-conditioned variances, hereafter denoted by *a*(*m*) for mone *m*. Similar to many ANOVA-based analyses this ratio measures distributional differences and is closely related to Bhattacharyya distance (see [42]). It converges to 1 for non-marker mones and converges to larger values for markers. Larger *a*(*m*) values denote larger distributional differences. Assuming balanced samples of sizes 200 and 100 we compute the *a*(*m*) values resulting in a posterior of 0.95 as thresholds to distinguish moderate [a(m)=1.088] and strong [a(m)=1.159] mones separating tumor from normal slides (see Supplementary Figure 2).

Structured multi-class OBF (see [43] for details) considers the four possible relations (known as structures by OBF) between frozen normal, frozen tumor, and FFPE tumor slides: (A) a mone does not differentiate between slides (prior probability=0.5), (B) a mone has one distribution for frozen slides (both tumor and adjacent normal slides) and another distribution for FFPE slides (prior probability=0.5/3), (C) a mone has one distribution for tumor slides (both frozen and FFPE) and another distribution for frozen normal slides (prior probability=0.5/3), and (D) a mone has one distribution for FFPE tumor slides and frozen adjacent normal slides and another distribution for frozen tumor slides. Mones with structure B for which frozen tumor and FFPE tumors lie on both sides of frozen adjacent normal slides (based on mean values) are considered ineffective due to FFPE/frozen differences.

### 4.3. Mone correlation analysis

For each cancer type, we calculated correlations between mones over all samples of the given cancer type. This analysis was done for each cancer type. We use the Ledoit-Wolf shrinkage [44] implementation of the scikit-learn python package [45] for computing covariance matrices. We then compute the correlation matrix from the covariance matrix. We apply the Fisher transform to correlation coefficients and approximate the null with its asymptotic Gaussian distribution. Benjamini-Hochberg [38] procedure is used for FDR correction. We use seaborn package[46] with default values to generate clustergrams. Correlation matrices are averaged to compute the pooled correlation matrix. Statistically significant mone-mone correlations are referred to as “correlated mone pairs.”

Differential mone correlations denote the difference between the correlation coefficient of tumor and normal slides, i.e., for each cancer, differential matrix is computed by subtracting the correlation coefficient matrix of normal slides form the correlation coefficient matrix of tumor slides. Differential mone correlation analysis uses the asymptotic Gaussian distribution of the difference between Fisher transformed correlation coefficients to compute the p-values. Statistically significant differential mone correlations are referred to as “differentially correlated mone pairs.”

### 4.4. Linear classification models

We implement MLDA and LR-LASSO using the scikit-learn python package [45] with default values except we set *C* = 100 and use the “saga” solver [47] in non-binary problems for LR-LASSO. We observed little sensitivity of AUCs to the *C* values ranging from 1 to 1000, and hence use *C* = 100 throughout. We randomly split data to train and test sets 10 times, which are then used to compute the mean and variance of the AUCs. We use the Scipy package [39] to implement the Wilcoxon signed rank test.

### 4.5. Cell segmentation and classification

Cellpose [48] was used to segment and count number of cells in BRCA, LUAD, and COAD tiles. Given the fixed magnification and tile size the number of cells per tile captures tile level cell density. Median number of cells per tile was used as slide level cell density index. HoverNet [49] was used to segment, count, and classify nuclei within COAD tumor slides. HoverNet was executed using the pre-trained PanNuke model ([50]), such that nuclei were classified into one of five types: neoplastic epithelial, non-neoplastic epithelial, connective (including fibroblasts and endothelial), inflammatory (including leukocytes, lymphocytes, and macrophages), and dead nuclei. Median number of cells nuclei across tiles were used as cell density. Median number of predicted inflammatory nuclei across tiles were used to characterize presence of immune cells.

#### 4.5.1. Integrative mone-gene analysis

Gene expression data were downloaded from the GDC portal [51]. We only used slide-gene expression pairs where both the slide and the expression profile were from the same vial. Log normalized FPKMs were used. Genes with zero counts in more than half the mone-gene pairs or expression standard deviation below 0.25 were removed. Given a set of mone-gene pairs, we stack the mone and gene vectors and compute the covariance matrix using the Ledoit-Wolf shrinkage method [44] implemented in the scikit-learn python package [45]. Correlation values are computed given the covariance matrix similar to mone correlation analyses. Statistical significance tests are performed similar to mone correlation analyses.

#### 4.5.2. Immune profiling and analysis

Leukocyte fractions of TCGA samples were obtained from [29]. All T-cell and B-cell categories were summed to obtain T-cell and B-cell proportions, respectively. The fractions of T-cell and B-cells were summed to obtain lymphocyte fractions. Log normalization of fractions were used throughout. Correlation analysis of immune scores with mones and IG score was performed similar to mone correlation analysis. B-cell percentages above 3% were removed for computing the B-cell correlations as they were deemed outliers.

## Acknowledgements

The authors would like to thank Jill Rubinstein for her feedback. A. F. would like to thank The Jackson Laboratory for the JAX scholar award.

## Conflict of interest

The authors declare no conflict of interest.

## Notes

https://portal.gdc.cancer.gov/

### Competing Interest Statement

The authors have declared no competing interest.

