## Supplementary Figures for "Deep Learning Features Encode Interpretable Morphologies within Histological Images"

By Ali Foroughi pour[1] and Brian White[1] and Jonghanne Park[1] and Todd B. Sheridan[1,2] and Jeffrey H. Chuang[1,3]

[1]*The Jackson Laboratory for Genomic Medicine, 10 Discovery Dr. Farmington, CT, 06032* ** **

[2]*Department of Pathology, Hartford hospital, 80 Seymour St, Hartford, CT, 06106*

[3]*Department of Genetics and Genome Sciences, UCONN Health, Farmington, CT, 06032*

### Supplementary Figures.

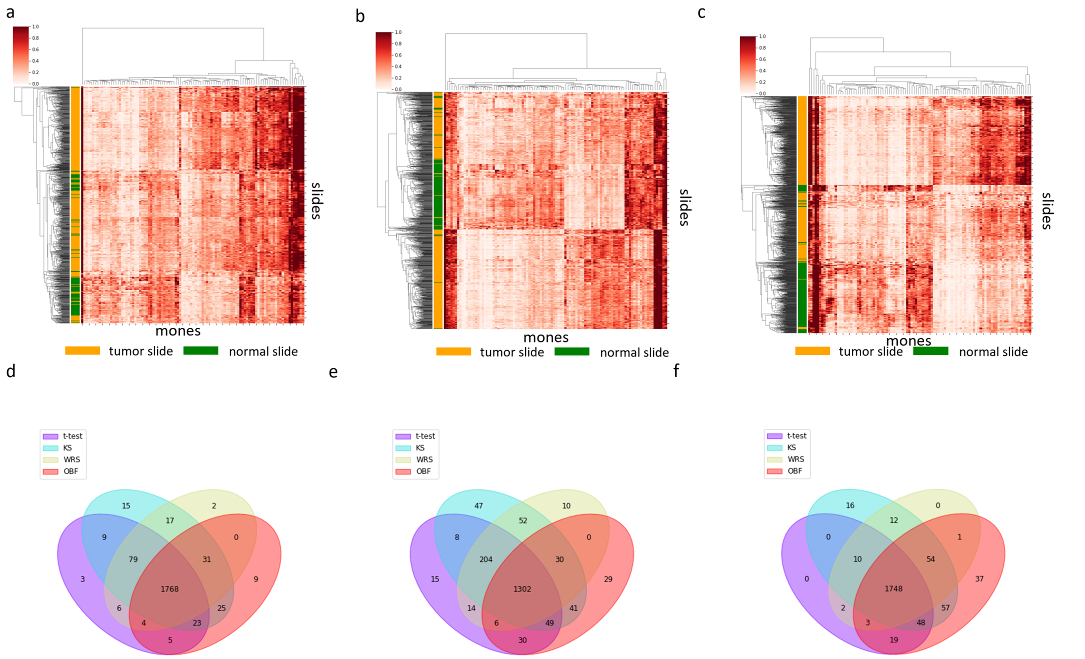


**Supplementary Figure** 1: **Mones strongly differentiate phenotypes.** Clustermaps of frozen adjacent normal and frozen tumors of (a) LUAD, (b) LUSC, and (c) KIRC using the top 100 OBF mones. Venn diagrams of the mones with distributional differences between tumor and normal slides according to different statistical tests in (d) LUAD, (e) LUSC, and (f) KIRC.

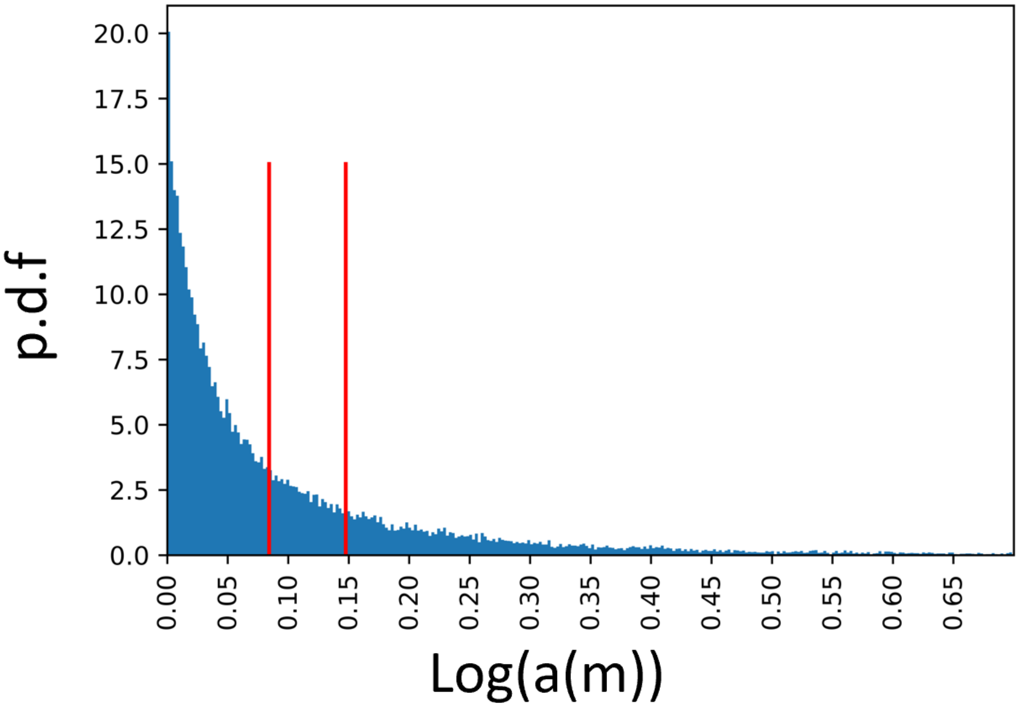


**Supplementary Figure** 2: **Mones show a wide range of distributional differences separating tumor and adjacent normal frozen slides.** Histogram of the extent of distributional difference between tumor and normal slides of all 19 cancers: $63\%$ of mones are weak or non-markers, $15\%$ are moderate tumor markers ($\log(a(m)) = 0.084$), and $22\%$ are strong tumor markers (log(a(m)=0.148) in differentiating tumor and normal slides. Larger $a(m)$ values denote larger distributional differences (see Methods).

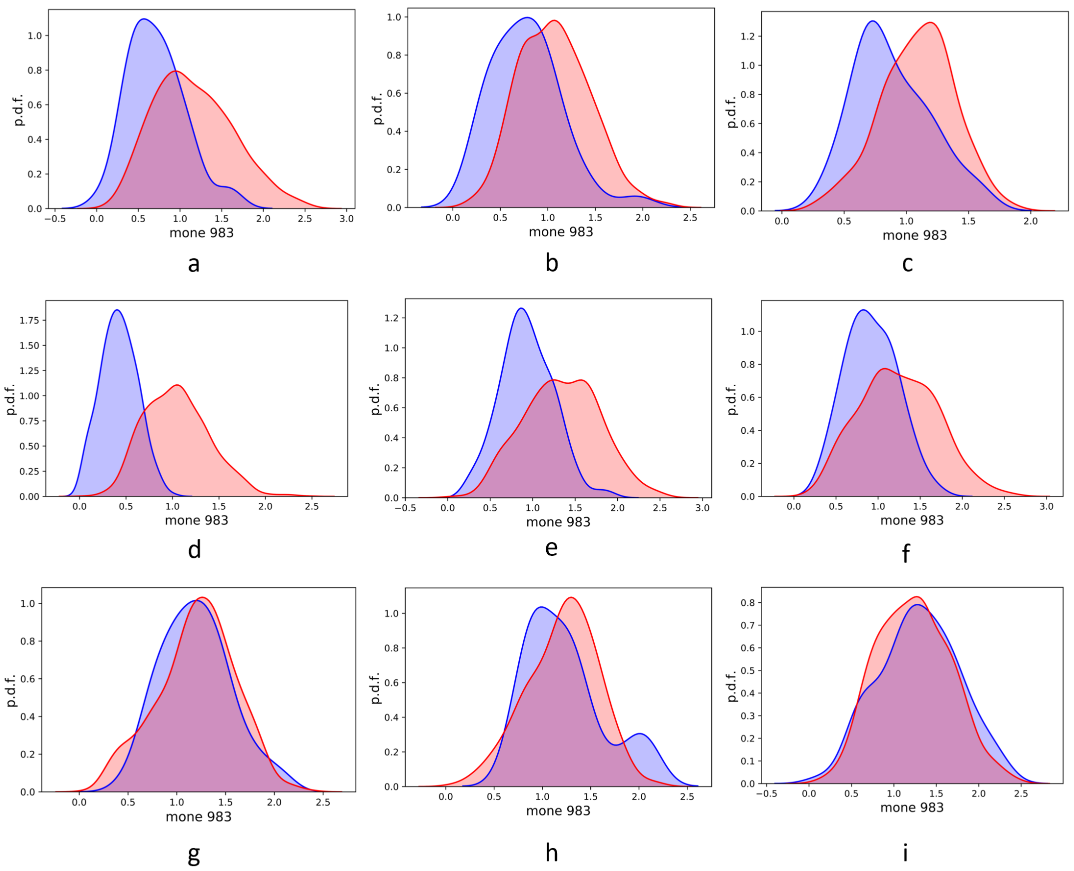


**Supplementary Figure** 3: **Mone 983 separates tumor and normal slides in several cancers.** Distribution of mone 983 between tumor (red) and normal (blue) frozen slides of multiple cancers: (a) HNSC, (b) BLCA, (c) PAAD, (d) BRCA, (e) LUAD, (f) LUSC, (g) COAD, (h) READ, and (i) STAD. All raw p-values in (a) HNSC, (b) BLCA, (c) PAAD, (d) BRCA, (e) LUAD, and (f) LUSC are $< 1e-3$. Mone 983 is not a tumor marker in (g) COAD, (h) READ, and (i) STAD.

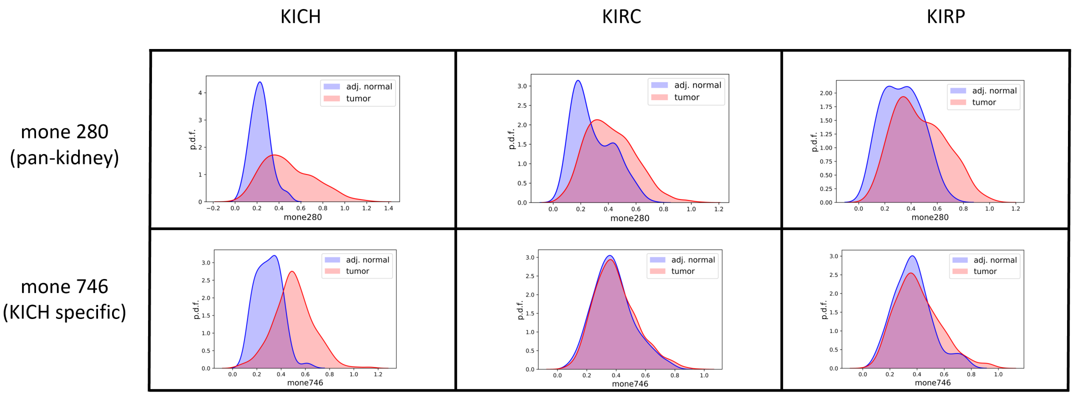


**Supplementary Figure** 4: **Mones may be cancer specific or show similar patterns across multiple cancers.** Example of a pan-kidney mone separating tumor and normal slides in all kidney cancers (Top). Example of a cancer specific mone only distinguishing tumor and normal slides of kich (Bottom).

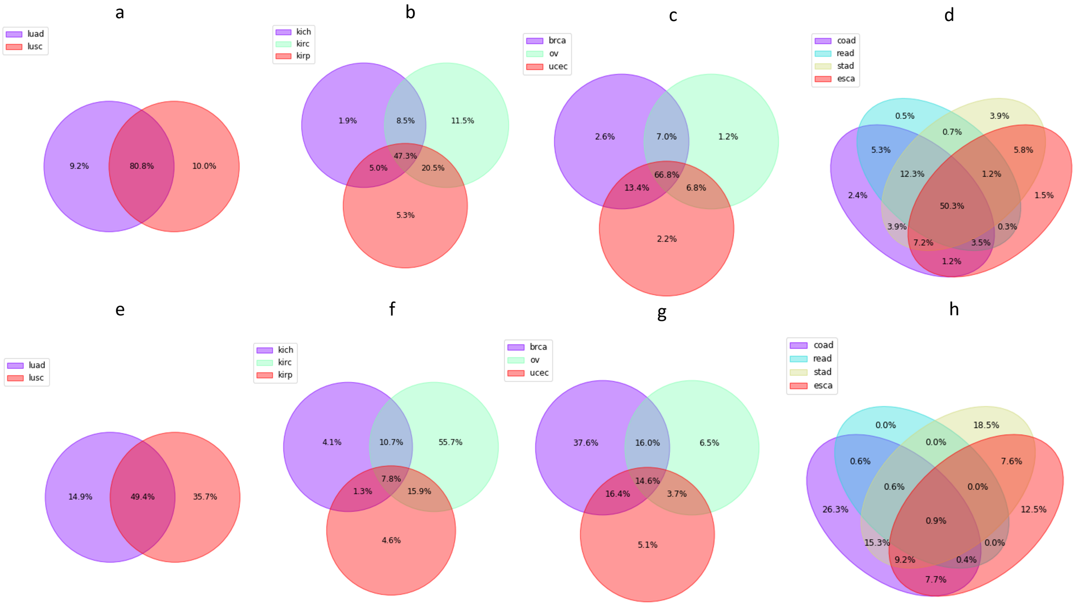


**Supplementary Figure** 5: **Mone correlations are conserved across tumors slides, but differential correlations are cancer specific.** Venn diagrams of percentage of correlated mone pairs across tumor slides of multiple cancer families (a)-(d). Venn diagrams of percentage of differentially correlated mone pairs (see methods) across tumor and normal slides of multiple cancer families (e)-(h). In all figures only mone pairs that were statistically significant in at least one of the cancers of the family are considered to compute percentages.

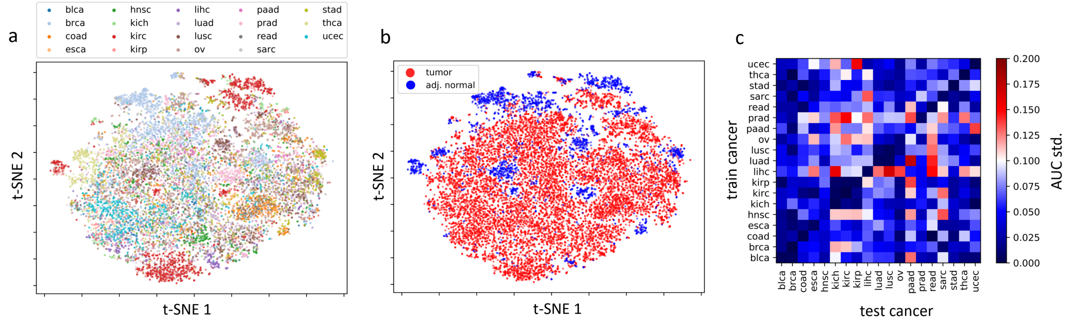


**Supplementary Figure** 6: **Mones separate frozen slides.** 2D t-SNE plots of tumor and normal slides of all 19 cancers using all 2048 mones colored based on (a) cancer type and (b) tumor/normal status. (c) standard deviation of the tumor/normal cross-classification AUCs using a mone based LR classifier.

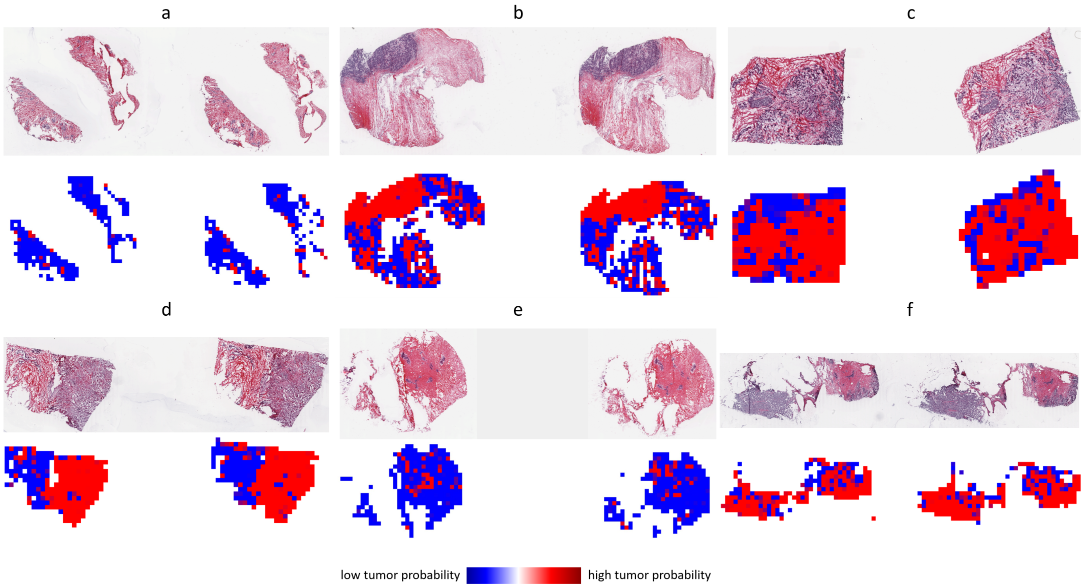


**Supplementary Figure** 7: **Slide level mone-based tumor/normal classifiers produce meaningful tile-level predictions.** Heat maps of tumor probabilities across tiles of several brca test slides. Note the LR-LASSO classifier is trained using slide level mones of the training set. (a) and (e) are adjacent normal slides. The rest are tumor slides.

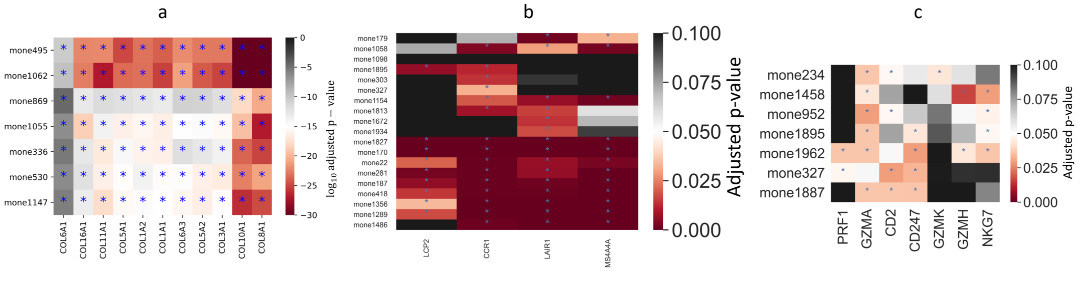


**Supplementary Figure** 8: **Adjusted p-values of mone-gene correlation analyses.** (a) log10 transformed adjusted p-values of the unsupervised mone-gene correlation analysis in OV. Adjusted p-values of correlation analyses of mones and immune genes in (b) pan-GI cancers and (c) LUAD. Starred adjusted p-values are statistically significant (FDR=5%)

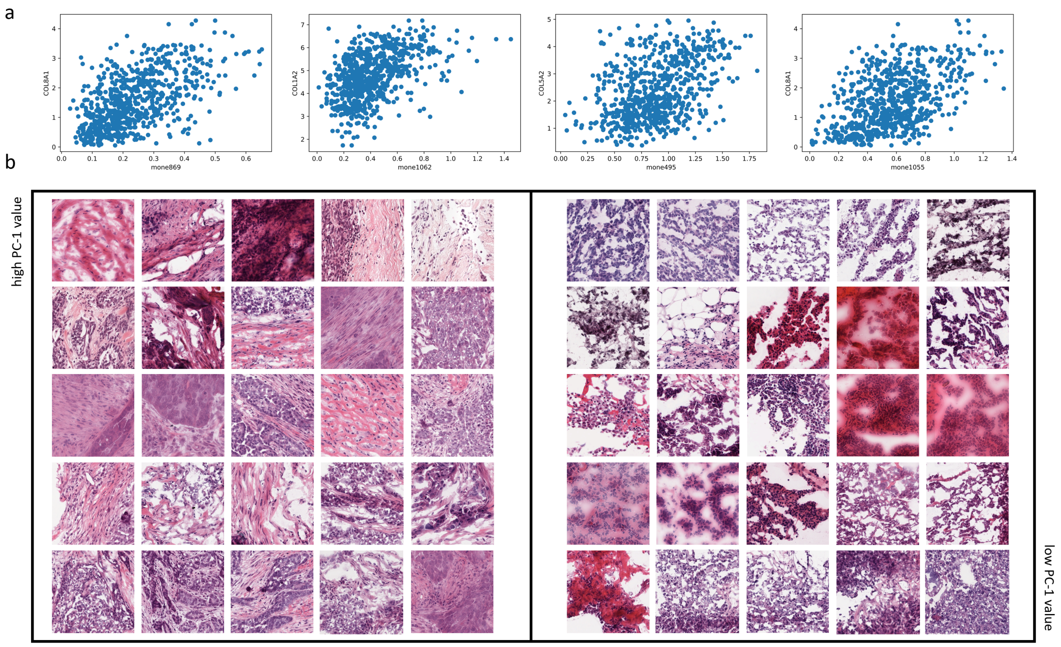


**Supplementary Figure** 9: **Mones encode collagen content in OV.** (a) Scatter plots of several correlated mone-gene pairs in OV. (b) Example tiles of slides with extreme (high or low) PC-1 values.

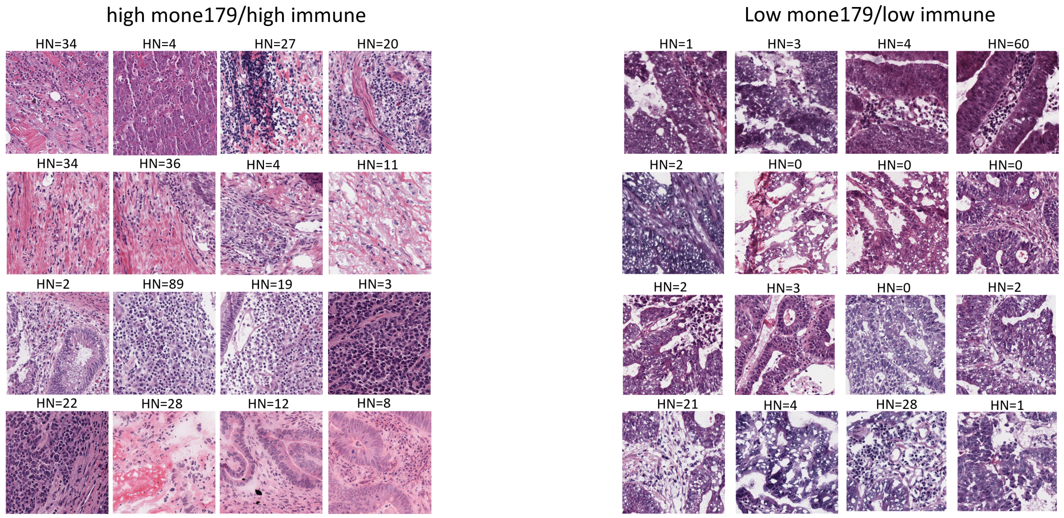


**Supplementary Figure** 10: **Mone 179 separates COAD slides based on immune infiltration.** Example coad tiles with exteme (high and low) values for mone 179. Given the positive correlation of mone 179 with leukocyte fraction and HoverNet scores high mone values should depict regions with high immune activity and vice versa. This correlation was verified by pathologist review. The numbers on top of each tile denote the number of immune cells in the tile according to HoverNet.

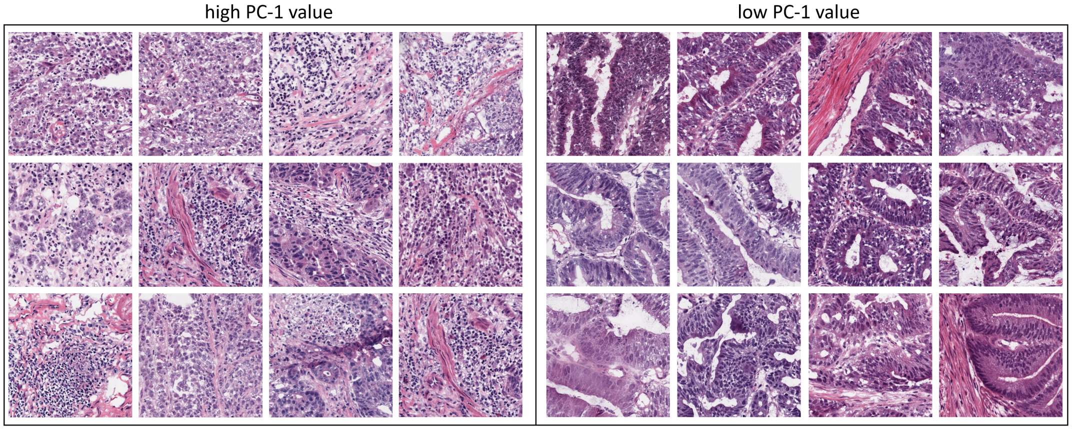


**Supplementary Figure** 11: **PC1 of GI immune mones separates COAD slides based on immune infiltration.** Example coad tiles with exteme (high and low) values for PC1 dimension of immune mones. Given the positive correlation of PC1 with leukocyte fraction and HoverNet scores high PC1 values should depict regions with high immune activity and vice versa. This correlation was verified by pathologist review.

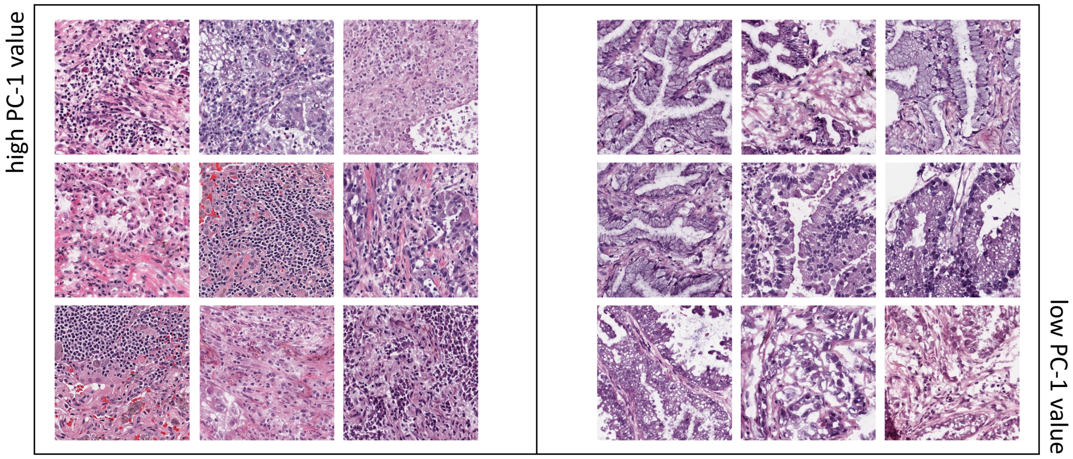


**Supplementary Figure** 12: **PC1 of LUAD immune mones separates slides based on immune infiltration.** Example LUAD tiles with extreme (high and low) values for PC1 dimension of immune mones. Given the positive correlation of PC1 with leukocyte fraction high PC1 values should depict regions with high immune activity and vice versa. Pathologist review verifies this correlation.
